# Analysis of Kinetic of eyelids of little owl *Athene noctua*

**DOI:** 10.1101/2022.08.20.504636

**Authors:** Fatma Abdel-Regal Mahmoud, Nahed Ahmed Shawki, Amany Mohamed Abdel-Mageed, Fatma A. Al-Nefeiy

**Affiliations:** Assiut University, Faculty of Science. Zoology Department, Assiut, Egypt; University of Jeddah, College of Science, Department of Biology, Jeddah, Saudi Arabia

**Keywords:** kinetic, eyelids, periorbital sheet, anatomy, little owl

## Abstract

The present study used the video’s recording technique to record the kinetic activity of lids, beside the anatomical, and the histological studies of the eyelids structure. The present authors found that the fundemental eyelids movements can be uniquely and reliably characterized by their anatomical relationships that confirmed also via video record of kinetic performance of these eyelids. The results show that levator palpebrae muscle split into many directions as a main generate motor to increasing the movement of the upper lid. In other side, the contraction of Depressor palpebral inferioris muscle together with the active upward forces of levator palpebrae muscle resultin g in opening the eye. While, the closure of lids arise from the passive downward forces and relaxation of the levator palpebrae and Depressor palpebral inferioris muscle as well as, the cooperation action of retractor anguli oculi lateralis and medialis muscle. The results also recorded that NM movement is reversely proportion to the level of kinetic of other eyelids.. The mobility of Nm in little owl occurs under effect artificially external stress. These anatomical data and sequence video records have confirmed that the upper eyelid move more compare to other lids. The present authors also suggest that the mobility of lids may stimulate through external pressure force of some surrounding structure like periorbital sheet. Also, the histological study exhibited that the structure of two eyelids is very similar in the little owl and the variability are showing in the number of cell layers that forms their epithelium of skin and palpebral surfaces, the distribution of pigment granules within them, as well as, the degree of keratinization on their surface.

**Summary:** This study gives a comprehensive description of eyelids movement in little owl and discusses the impact of some surrounding conditions in kinetic performance of these eyelids

## Introduction

Owls have a different activity patterns, from nocturnal to crepuscular or cathemeral, or to diurnal. The little owl is a predator bird, hunting mainly at night but also active during daylight (Zerunian et al., 1982). In 2008, Hall classified the little owl as phototropic-birds (diurnal species). Obviously, the activity pattern of the little owl is still under debate and needed to more study to confirm it. The difference of activity pattern is reflected in the structure of visual system of owls (Gutierrez-ibariez et al., 2012). Several previous studies have mentioned the structure of the visual system of some other owls’ species but we noticed that little owl was not given much of such attention.

The protective apparatus of eye (ocular adnexa) includes lids, glands and muscles, this apparatus revealed significance differences in bird, reptiles, in mammals that reflect a wide range of the activity patterns (Schramm et al 1994; Klec’kowska-Nawrot et al., 2016, 2017). The three eyelids are a common feature in bird, reptiles and in some mammals (Jochems and Philips, 2015). Many previous studies have concluded that the upper eyelid in different bird species is thick and short while the lower eyelid is thinner, longer, and moveable (Hall et al., 2009 ; Jochems and Philips, 2015). They were also recorded that the third eyelid is elastic membrane and moves rapidly (Montiania-Ferreira, 2001). The main purpose of this apparatus is to cleanse and lubricate the front of the cornea, as well as, it act as an immune-protective for the eye (Klec’kowska-Nawrot et al., 2017).

Klec’kowska-Nawrot et al. (2017) recorded wide variations in the microstructure of eyelids in many wild bird species. These recent studies confirmed that the eyelids of bird differ in their structure among different avian species. for this reason, the present study attempts to give more details about structure of eyelids of some birds had been neglect in this study field, like the little owl (*Athene noctua*) and discuss the influence of some conditions in kinetic performance these eyelids.

## Material and methods

Ten adult samples of little owl were brought to the comparative anatomy of vertebrate laboratory, Zoology Department, Faculty of Science, Assiut University, in good health condition and care them according to the guidelines of the research ethics committees, Assiut University (www.enrec.org). The samples of little owl placed in a wide cage and a camera was installed in one of the corners of the cage for a period of 24 hours to follow the natural movement of the eyelids without any external influence. In the next day, the samples were caught and the eyelids (upper and lower) were fixed using the fingers of the hand and the resulting movement was recorded. The ten samples were divided into the following procedures; seven samples to conduct the anatomical study and prepare the skull, and three samples for histological study. For anatomy study, the head was fixed in 10% formalin for two weeks and then store in 2% phenoxy-ethanol for long term preservation. The photos were taken by Toup camp XCAM full HD camera. The present study are followed Nomina Anatomica Veterinaria (2017) for anatomical terminology of the eyelid’s musculature system and cranial skeleton.

For light microscopic investigations, Specimens of eyelids were fixed in 10% neutral formalin for three days then put in 70% ethyl alcohol for two days, after which were dehydrated through a series of ethyl alcohols, cleared in methyl benzoate for three days, embedded in paraffin wax, sectioned serially (7μm) and then sections stained with Haematoxylin and Eosin, Masson’ trichromic stain, and periodic acid-Schiff (PAS) (Drury and Wallington, 1980). for Transmission Electron Microscopic investigation (TEM), parts from the eyelids were cut into small pieces (1mm each) and were fixed in a cacodylate-buffered-solution of 5% glutraldehyde for 2-hrs, then washed several times in the same buffer for 1-hr at pH=7.2, then the specimens were post fixed in a cacodylate buffer 1% Osmium tetraoxide for 2-hrs at 4oC. Specimens were washed several times as in the second step, and then it was followed by dehydration in graded series of alcohol. The specimens were embedded in epoxy resin; they were treated for semi-thin sectioning at 1m thickness and stained with toludine blue for light microscopic examination. The photos were taken by Olympus camera model DP74 connected with Olympus microscope model BX43.

For scanning electron microscopy, eye were cut to small pieces and directly fixed in 5% glutaraldehyde in a cacodylate buffer for 48 hr at 4ºC and washed in three changes of 0.1 % cacodylate buffer, then the specimens were post-fixed in a cacodylate buffered solution of 1% osmium tetroxide for 2 hr at 37ºC. The specimens were washed in the same buffer three times, dehydrated through ascending series of ethyle alcohol and then infiltrated with amyl acetate for two days. The drying of specimens is accomplished by the critical point drying using liquid carbon dioxide, mounted and sputter-coated with gold. The specimens were examined on a Jeol scanning electron microscope (J S M-5400I V), at 15 kv.

## Results

The eyelids of the little owl *Athene noctua*, distinguished into; upper, lower and third eyelids

### The anatomical investigation of the upper and lower eyelids of little owl

The little owl (*Athene noctua*) has movable eyelids (upper eyelid, *palpebral dorsalis* and lower eyelid, *palpebral ventralis*). The upper eyelid is shorter, thicker and high movable more than the lower one (Figs.6& 8). The palpebral margin “*plica marginalis*” of each eyelid is deeply pigmented and segmented into number of folds. The anatomical investigation of the eyelids of the little owl (*Athene noctua*) reveals that the anterior edge of palpebral margin of upper lid is raised into voluminous ridges which give sausage-like segments. These segments decrease in size posteriorly and convert into shallow irregular ridges (Fig.6b). While the palpebral margin of lower eyelid bears relatively larger anterior segment than that of its posterior edge (Fig.6a). Moreover, the palpebral margin of each eyelid carries two rows of modified feathers as eyelashes (Fig. 6a& b).

### Histological and scanning electron microscopical investigations of the upper eyelid

Histological investigation of the upper eyelid of little owl (*Athene noctua*) showed that the skin surface of upper lid is composed of keratinized stratified squamous epithelium with three or four nucleated cell layers (Fig. 1a) which multiple into ten layers to form the palpebral margin of the upper eyelid of little owl (*Athene noctua*) lacking keratinization (Fig.1b). Numerous pigmented granules fill the cytoplasm of these cell layers of skin surface which increase in palpebral margin of upper eyelid (Fig.1b).This skin epithelium inclined inwards to form the conjunctival surface of upper eyelid which changes from stratified squamous type to stratified cuboidal (Fig.1c). The epithelium of conjunctival surface containing numerous goblet cells which exhibits purple color with PAS reaction and slightly bluish color with toluidine blue stain (Fig. 1c& d).

**Fig. 1.**
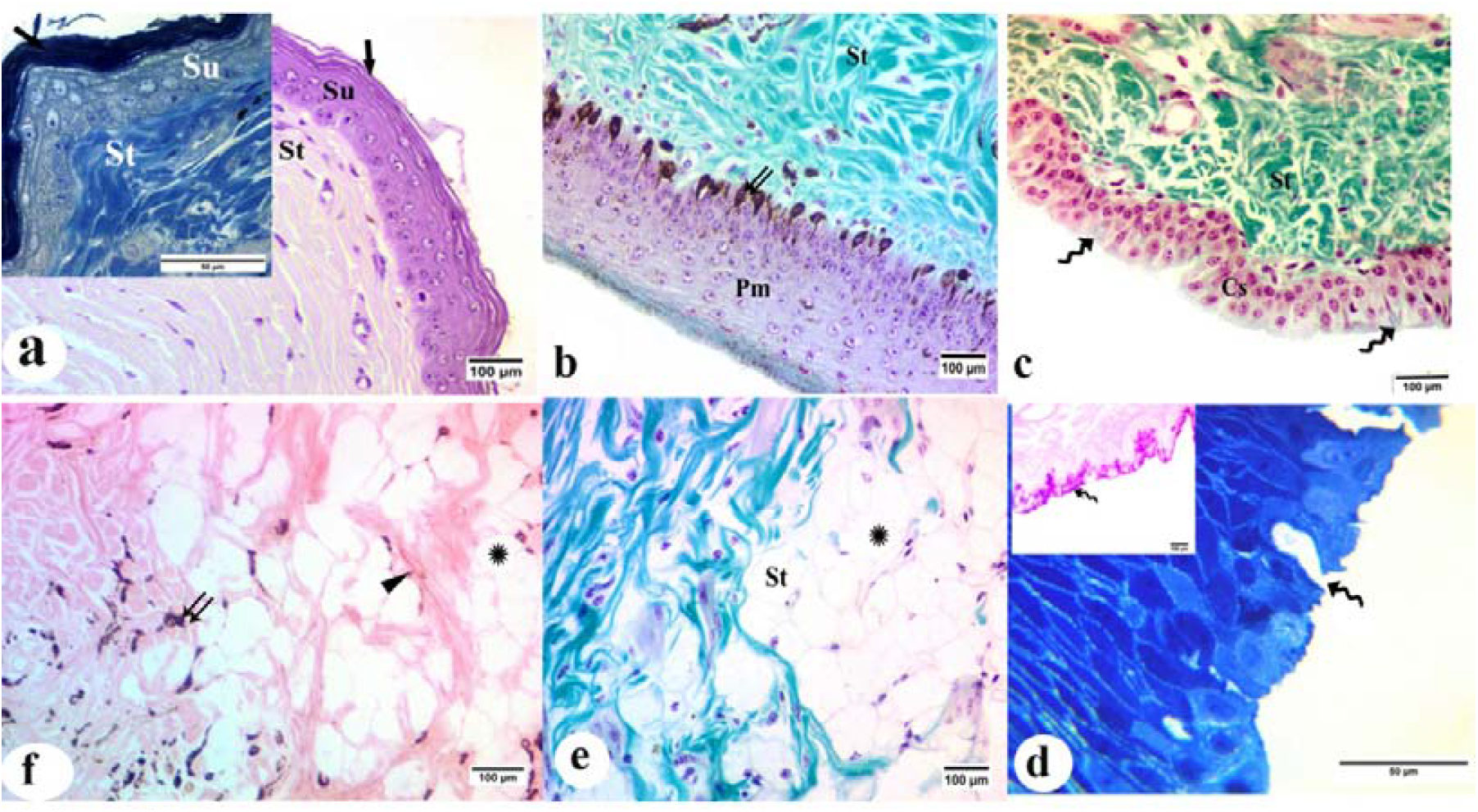
Photomicrograph of a transverse section of upper eyelid of little owl, *Athene noctua*, showing; (a) The main layers of skin surface (Su) is composed of stratified squamous epithelium covers with wavy keratin (arrow); upper left, show the parallel distribution of fibers beneath the surface (a by H& E, upper left by Toluidine blue, scale bar,100, 50µm). (b) The palpebral margin (Pm), numerous pigmented granules (double arrows), the collagen fibers within stroma (St) (Trichromic, scale bar, 100µm). (c& d) The conjunctival surface (Cs) with goblet gland (zigzag arrow) (c by Trichromic and d by Toluidine blue, upper left by PAS, scale bar, 100, 50µm). (e & f) Scattered network of fibers within stroma (St); collagen fibers (green color), elastic fibers (arrowhead) and clusters of adipocytes (star) (e by Trichromic, f by Orcien, scale bar, 100µm).

The stroma of upper eyelid contains scattered network of collagen fibers with less abundant elastic ones, sparse number of melanocytes and blood vessels. Also, Clusters of adipocytes fill the stroma of upper eyelid but decrease within its margin (Fig.1 e& f)

Semithin sections of the mucosa of skin surface of upper lid reveals the main following layers; stratum basal in which the basal cell layer containing rounded and oval nuclei; the stratum intermedium that forms two layers of polyhedral cells outer to the basal layer; the stratum transitivum that lies outer to the stratum intermedium and is formed of flattened cells with oval nuclei; and the stratum corneum is formed of wavy keratin which lacking nuclei (Fig.1 a).

Scanning electron microscopy (SEM) reveals the presence of heavy detached keratin on the skin surface of upper lid (Fig.3a). In addition, the appearance of goblet cells scattered on the conjunctiva surface of the upper eyelid with their secretion spreading on its surface. As well as, the intercellular borders are clear observed (Fig.3 b).

### Histological and scanning electron microscopical investigations of the lower eyelid

The structure of skin and conjunctival surfaces (Cs) of lower eyelid is similar to that of the upper eyelid. The skin surface of lower eyelid of little owl (*Athene noctua*) is composed of stratified squamous epithelium which consists of one or two nucleated cell layers and covers by thin keratin (Fig.2 a, b& c). These epithelial layers increase gradually inward the conjunctival surface to become ten non-keratinized nucleated cell layers, as well as, their cytoplasm contains highly dense of dark pigment granules especially at the palprebral margin (Fig.2 d).

**Fig. 2.**
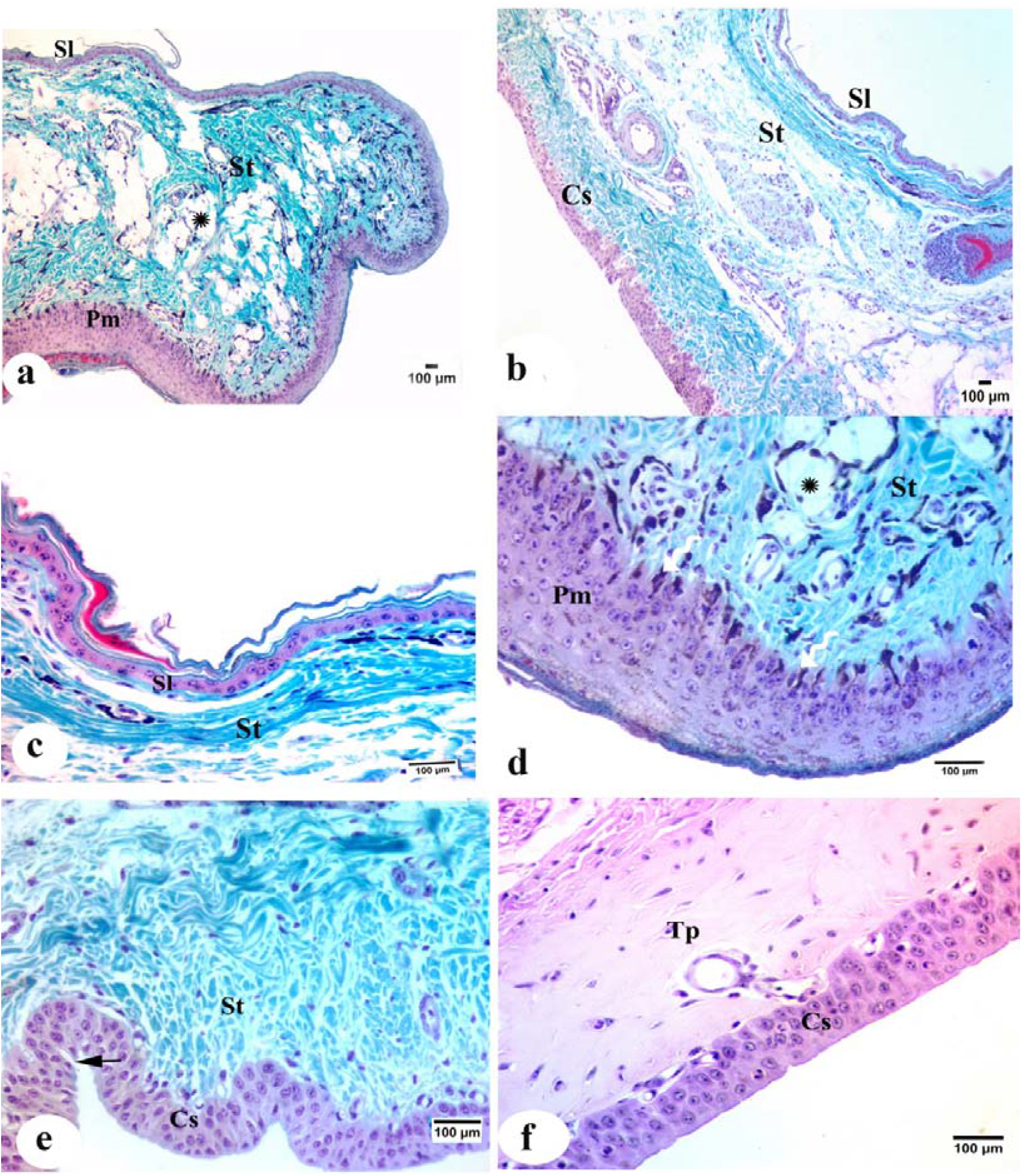
Photomicrograph of a transverse section of the lower eyelid of little owl, Athene noctua, showing; (a) the change in thickness of epithelium and keratinization along the lower eyelid, stroma (St) contains highly dense of dark pigment granules (zigzag arrow) especially at the palpebral margin (Pm) and adipocytes (star). (b) skin surface or lower eyelid (SI), stroma (St) with parallel collagen fibers. (c) The palpebral margin (Pm) highly dense of dark pigment granules (zigzag arrow) and adipocytes (star). (d) The conjunctival surface (Cs) with evidence goblet gland (arrow). (f) Inferior tarsal plate (Tp). (a, b, c, d, e by Trichromic, f by H&E, scale bar, 100µm).

**Fig. 3.**
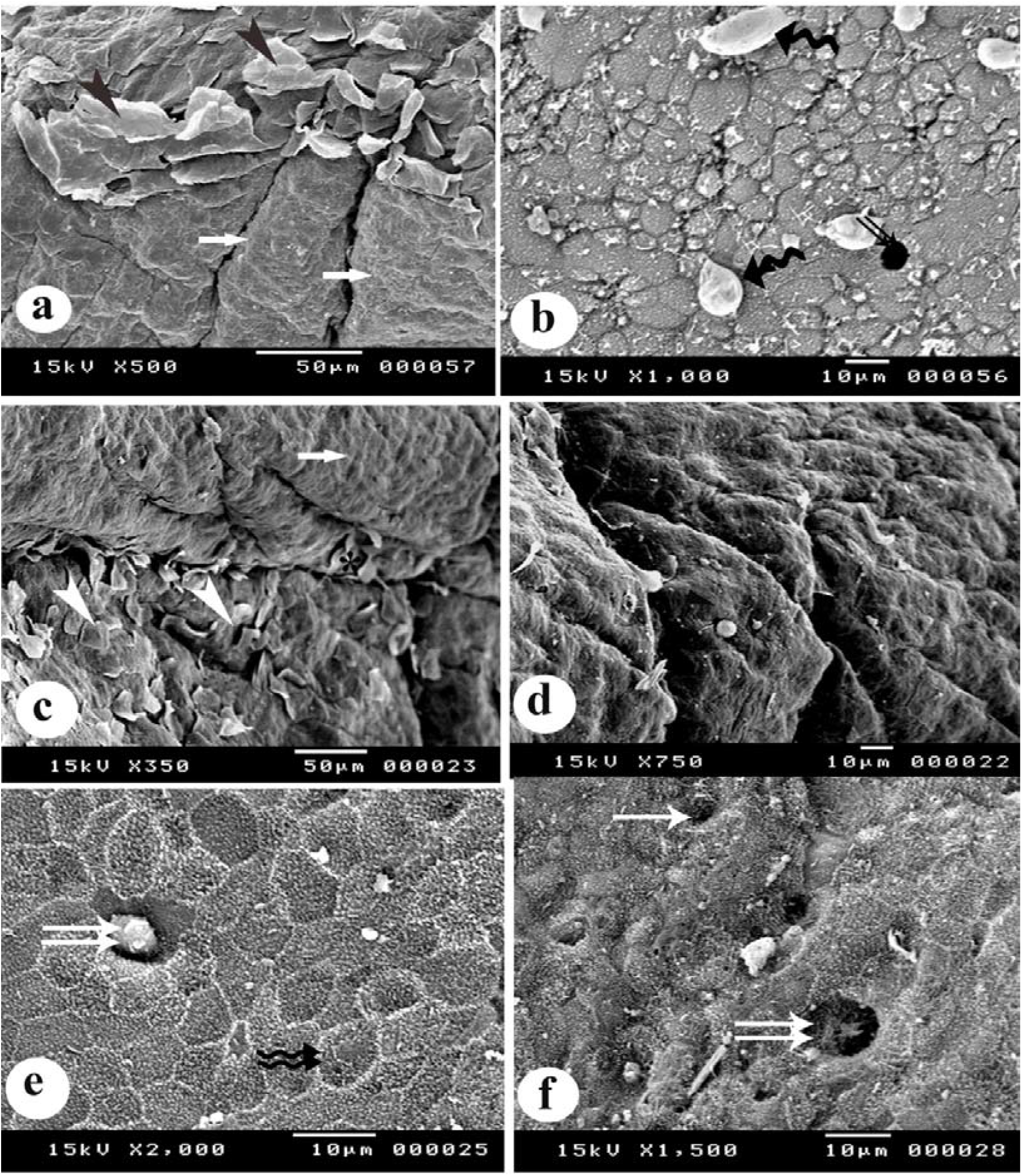
Scanning electron micrograph eyelids of the little owl, *Athene noctua*, showing; (a) the detached keratin in the skin surface of upper eyelid (arrowhead) which convert in smooth surface of the palpebral margin (arrow). (b) The conjunctival surface of the upper eyelid, pore of goblet cells (double arrows) and its secretion on surface (zigzag arrow). (c) The skin surface of lower eyelid with deciduous keratin (arrowhead) which disappears toward the conjunctival surface of the lower eyelid (arrow). (d) The smooth palpebral margin of lower eyelid. (e & f) the cell boarder of the conjunctival surface of lower eyelid (double zigzag arrows) and the presence of goblet cell (double arrows).

The epithelial layer that forms the conjunctival surface near the palprebral margin of lower eyelid, covers by thick keratinized layer. This thick keratinized layer return to disappear along the whole length of conjunctival surface (Fig.2 b, e &f).

The conjunctival surface of the lower eyelid of little owl (*Athene noctua*) is covered by stratified cuboidal epithelium with few goblet cells (Fig.2e). The plate of dense collagen fibers (inferior tarsal plate) is present beneath the conjunctival epithelium. That tarsal plate is invisible in the upper eyelid (Fig.2f).

The stroma of the lower eyelid is filled with parallel collagen fibers just underlies the skin surface of lower eyelid then become irregular distributed at the palpebral margin and beneath the conjunctival surface. It has been observed that numerous melanocytes, blood vessels and clusters of adipocytes are scattered within this stroma but the later one is less than that of the upper eyelid (Fig.2).

Scanning electron microscopy reveals the presence of heavy detached keratin on the skin surface of lower lid (Fig.2c &d), which disappears along the whole length of conjunctival surface (Fig. 2e& f). Moreover, the conjunctival surface of lower eyelid of little owl (*Athene noctua*) reveals appearance of goblet cells with their secretion spread on its surface and the intercellular borders are clear observed (Fig. 2e& f).

### The anatomical investigations of the third eyelid (nictitating membrane)

The nictitating membrane of little owl is well developed semi-transparent and mobile organ which formed as reduplication of conjunctiva (Fig.6c). That membrane possess two surfaces; the external surface which faces to the conjunctiva surface of upper eyelid (Palpreal surface) and merges superiorly with the fornix conjunctivae palprebrae, (Fig.4a) and internal one faces to the cornea (Bulbar surface) which merges with the fornix conjunctivae bulbi (Fig.4b). The free margin of this membrane is a thick and dark edge which is extended obliquely to the anterior canthus which is almost visible during rest of action beneath the palpebral margin of upper eyelid (Fig. 6a, b &c).

During observation of the little owl in laboratory, the movement of nictitating membrane of little owl (*Athene noctua*) showed that the membrane slips rapidly and obliquely across the front of eye from fixed point at the anterior canthus and the fornix of conjunctivae palprebrae towards ventrotemporal direction. This movement occurs when fixed the movement of eyelids. The anterior angle of membrane slips beneath the lower eyelid, additionally, the movement of both eyelids (Fig. 9)

### Histological and scanning electron microscopical investigations of the nictitating membrane

Histological investigation of the nictitating membrane of little owl (*Athene noctua*) reveals that both surfaces of nictitating membrane are covered by folded stratified epithelium (Fig.4a& b). The free margin and palpebral surface of the nictitating membrane are covered by non-keratinized squamous epithelium (Fig.4 c& e) while its bulbi surface is covered by stratified columnar epithelium with dome shape (Fig.4f). The epithelium of bulbi surface just near the free margin carries long cytoplasmic extension like the feather duster (Fig. 4d& f). Moreover, numerous of pigmented granules are interspersed only between the epithelial cells of free margin (Fig.4 c).

**Fig. 4.**
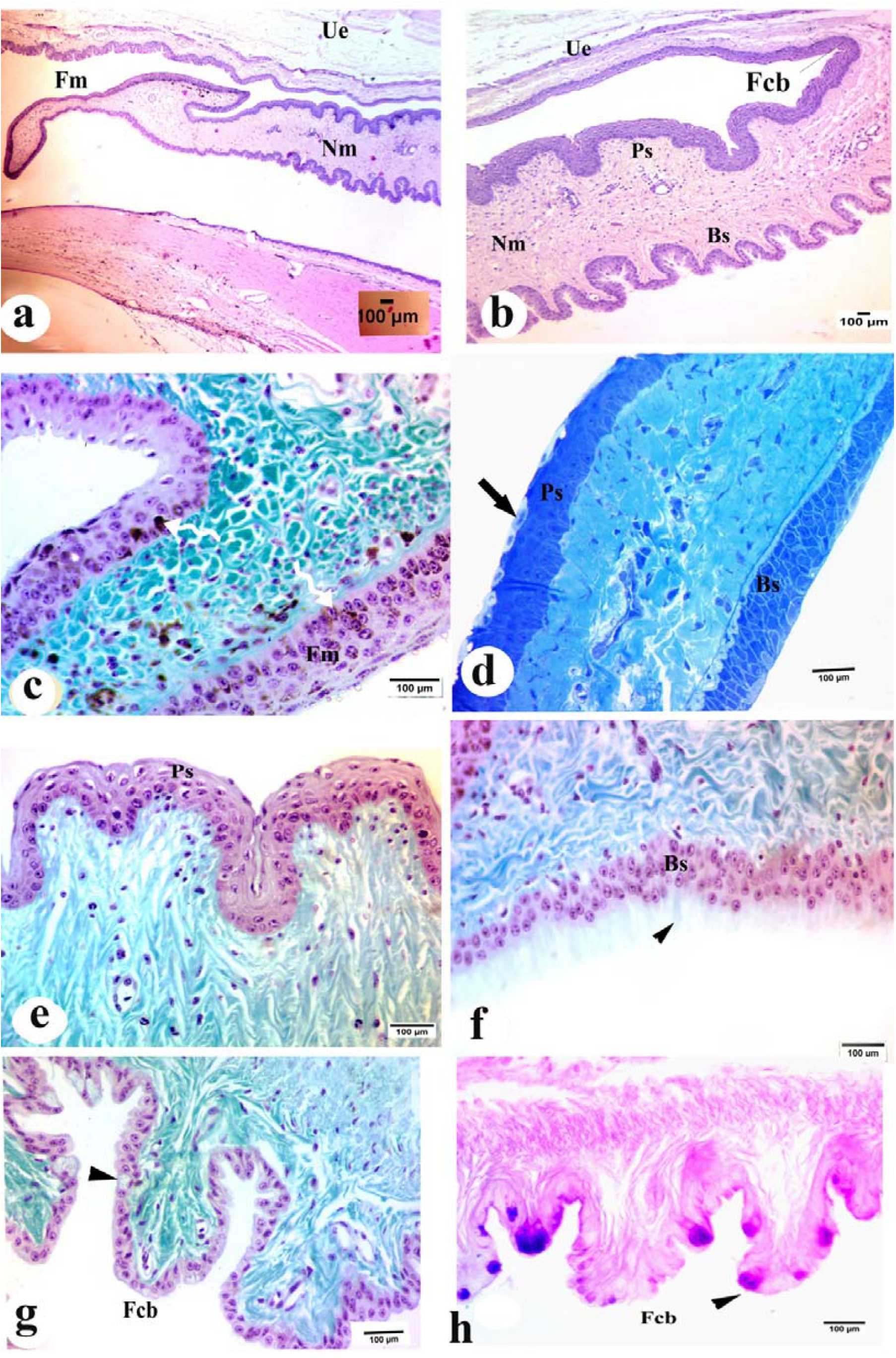
Photomicrograph of a transverse section of the nictitating membrane of little owl, *Athene noctua*, showing; (a) the surfaces of nictitating membrane; the palpreal surface (Ps), bulbar surface (Bs) and the free margin (Fm). (H& E, scale bar 100µm). (b) The Palpreal (Ps) and Bulbar surface (Bs) of nictitating membrane are covered by folded of stratified epithelium and fornix conjunctivae palprebrae (Fcb). (H& E, scale bar 100µm). (c) The free margin (Fm) of the nictitating membrane is covered by non-keratinized squamous epithelium, pigmented granules (zigzag arrow). (Trichromic, scale bar 100µm). (d) The palpreal surface (Ps) with superficial cells has pale cytoplasm (arrow), bulbar surface (Bs) (toluidine blue, scale bar 100µm). (e) The palpebral surface (Ps) of the nictitating membrane is covered by non-keratinized squamous epithelium (Trichromic, scale bar 100µm). (f) The bulbi surface (Bs) is covered by stratified columnar epithelium with long cytoplasmic extension like the feather duster (arrow head). (Trichromic, scale bar 100µm). (g& h) the epithelial of fornix conjunctiva bulbi (Fcb) which interspersed with numerous unicellular mucous glands (arrow head) which exhibits positive reaction with PAS (g by Trichromic, h by PAS, scale bar 100µm)

Semithin sections of the nictitating membrane of little owl (*Athene noctua*) reveal that the epithelium of the palpepral surface, the intermediate cells are flattening superficially to give squamous cell layers with oval nuclei and pale cytoplasm (Fig. 4d). On bulbi surface, the apical membrane of these intermediate cells is bulging to form dome-shaped cells with apical cytoplasm extension which are well observed by SEM (Fig.5c). That cytoplasm extension decreases in density toward the fornix conjunctiva bulbi (Fig. 5d), as well as, the epithelial of fornix conjunctiva bulbi (Fcb) is interspersed with numerous unicellular mucous glands which exhibits positive reaction with PAS (Fig.g& h).

**Fig. 5.**
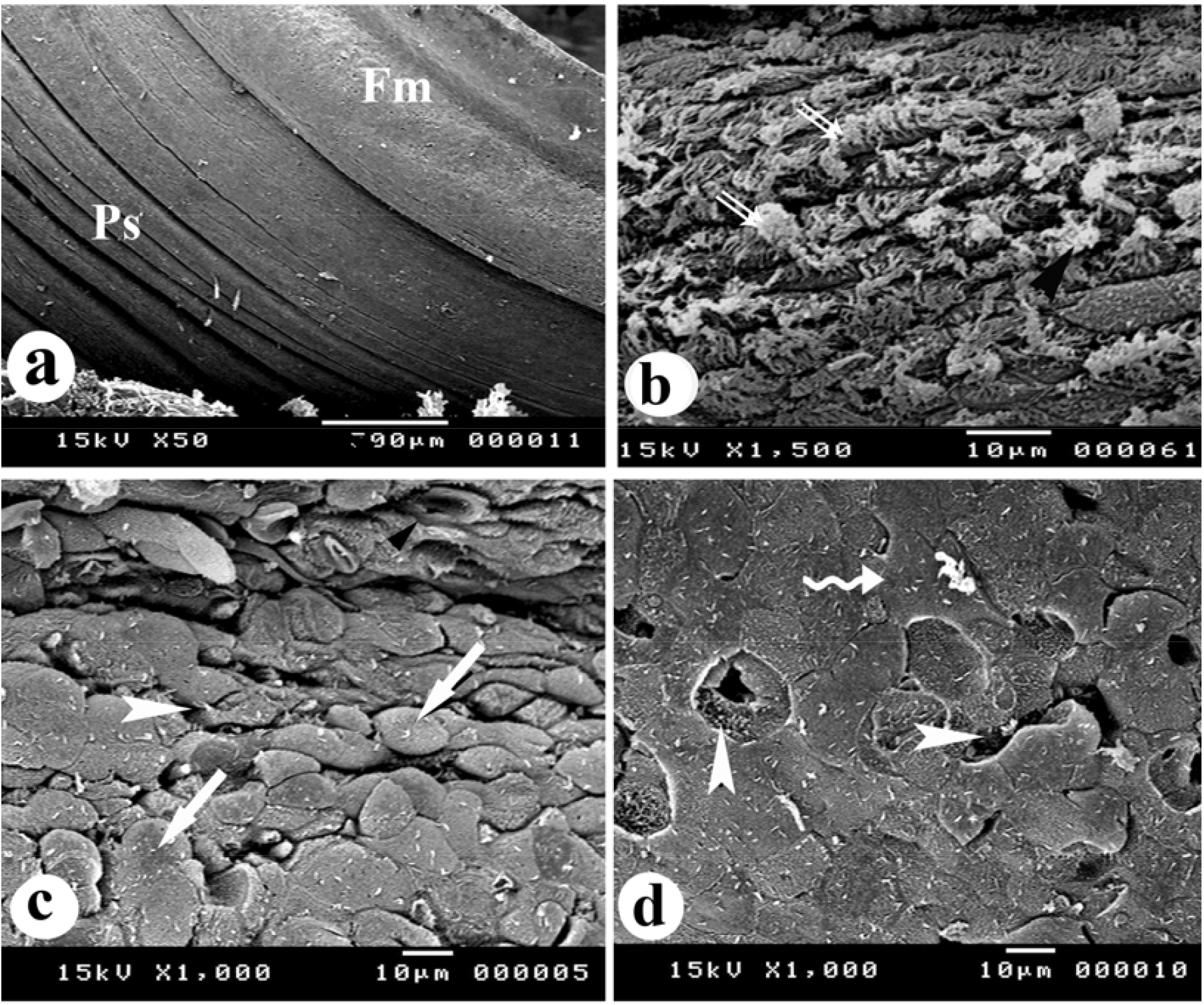
Scanning electron micrograph of the nictitating membrane of eye of little owl, Athene noctua, showing ; (a) the membrane becomes more wrinkle medially than its anterior and posterior fixed point at anterior and posterior canthus. (b) The long cytoplasmic extension which like the feather duster on the bulbi surface (double arrows). (c& d) The bulbi surface of the membrane have dome-shaped cells (arrow) becomes more flattened towards fornix conjunctiva (zigzag arrow), interspersed with scattered pores (arrow head.).

**Fig. 6.**
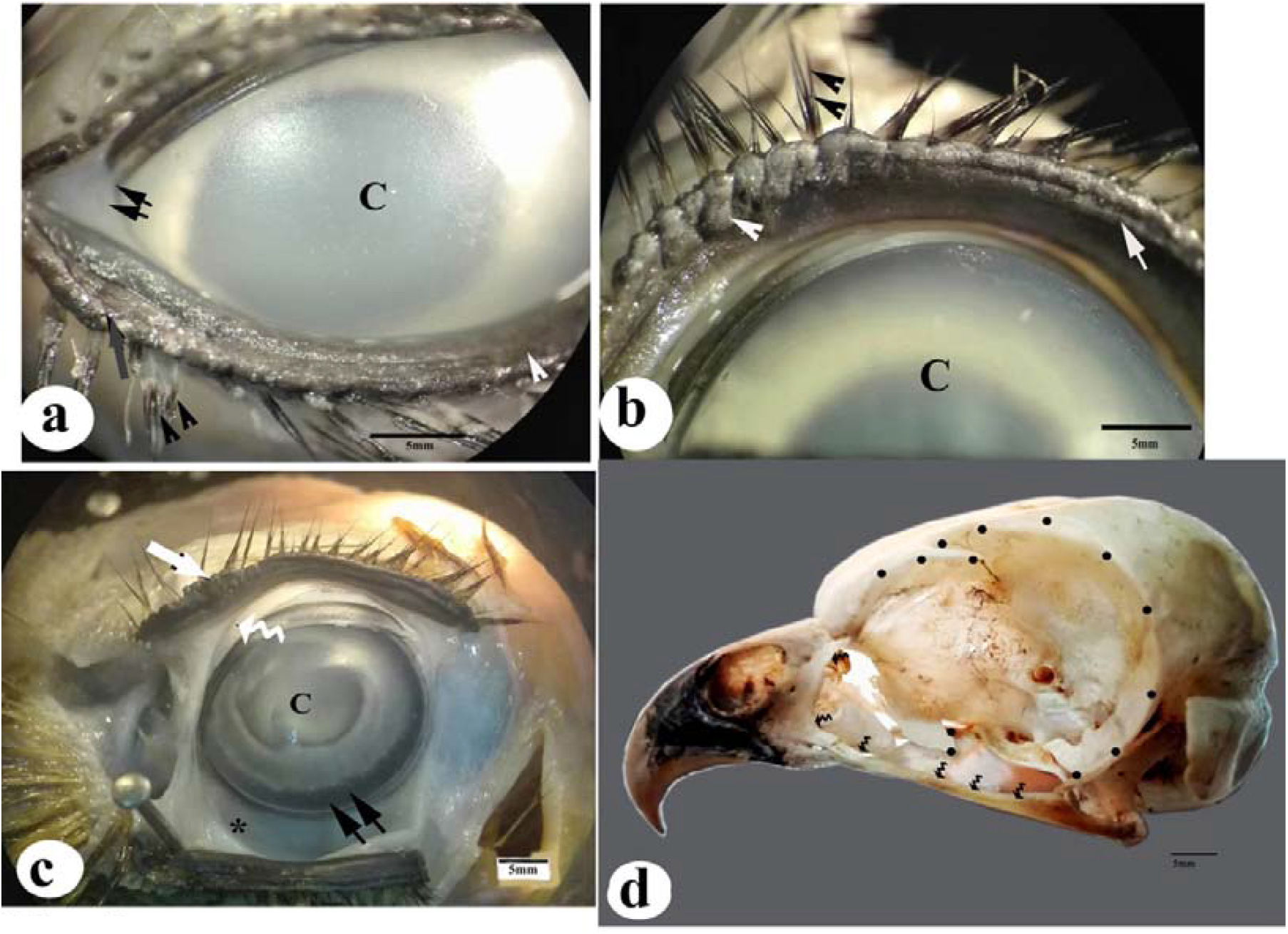
Photomacrograph of the eye of little owl, *Athene noctua*, showing; (a) the palpebral margin of the lower eyelid bears relatively large anterior segment (arrow) than that on it’s the posterior edge of palpebral margin (arrow head) and carried two rows of modified feathers as eyelashes (double arrows head), the fixed point of the nictitating membrane at anterior canthanus (double arrows). (b) the anterior edge of palpebral margin of the upper lid is raised into voluminous ridges which give sausage-like segments (arrow head) which decrease in size posteriorly and convert into shallow irregular ridges (arrow) and the eyelid carried two rows of modified feathers as eyelashes (double arrows head), the cornea (C). (c) The nictitating membrane is semi-transparent, formed as reduplication of conjunctiva (zigzag arrow), upper eyelid (white arrow), and lower eyelid (star), cornea (C), and irodocorneal junction (double arrows). (d) The attachment site of the periorbital sheet on the skull (dark spots) and indirectly attaches with some movement bony elements of skull (zigzag arrow).

Scanning electron microscopy (SEM) investigations of the nictitating membrane of little owl (*Athene noctua*) observes that the surface of this membrane becomes more wrinkle medially than its anterior and posterior fixed point (at anterior and posterior canthus) (Fig.5a). The bulbi surface of this membrane has dome-shaped cells interspersed with scattered pores (Fig.5c) and then this surface becomes more flattened towards fornix conjunctiva (Fig.5d).

### The Anatomical investigation of eyelids muscle (Fig. 7& Table 1)

These muscles control on the movement of the eyelids and nictitating membrane that including; muscle Levator palpebral Superioris (LPS), Levator anguli oculi (LAO), Retractor anguli oculi lateralis (RAOL), Retractor anguli oculi medialis (RAOM), Depressor palpebral inferioris (DPI), Quadratus (Qm),and Pyramidalis (Pm). All these muscles are covered completely by the periorbital sheet which a white strong sheet is appearing as parallel bundles of collagen fibers beneath the skin surface of eyelids (Figs. 1a& 2c)

**Fig. 7.**
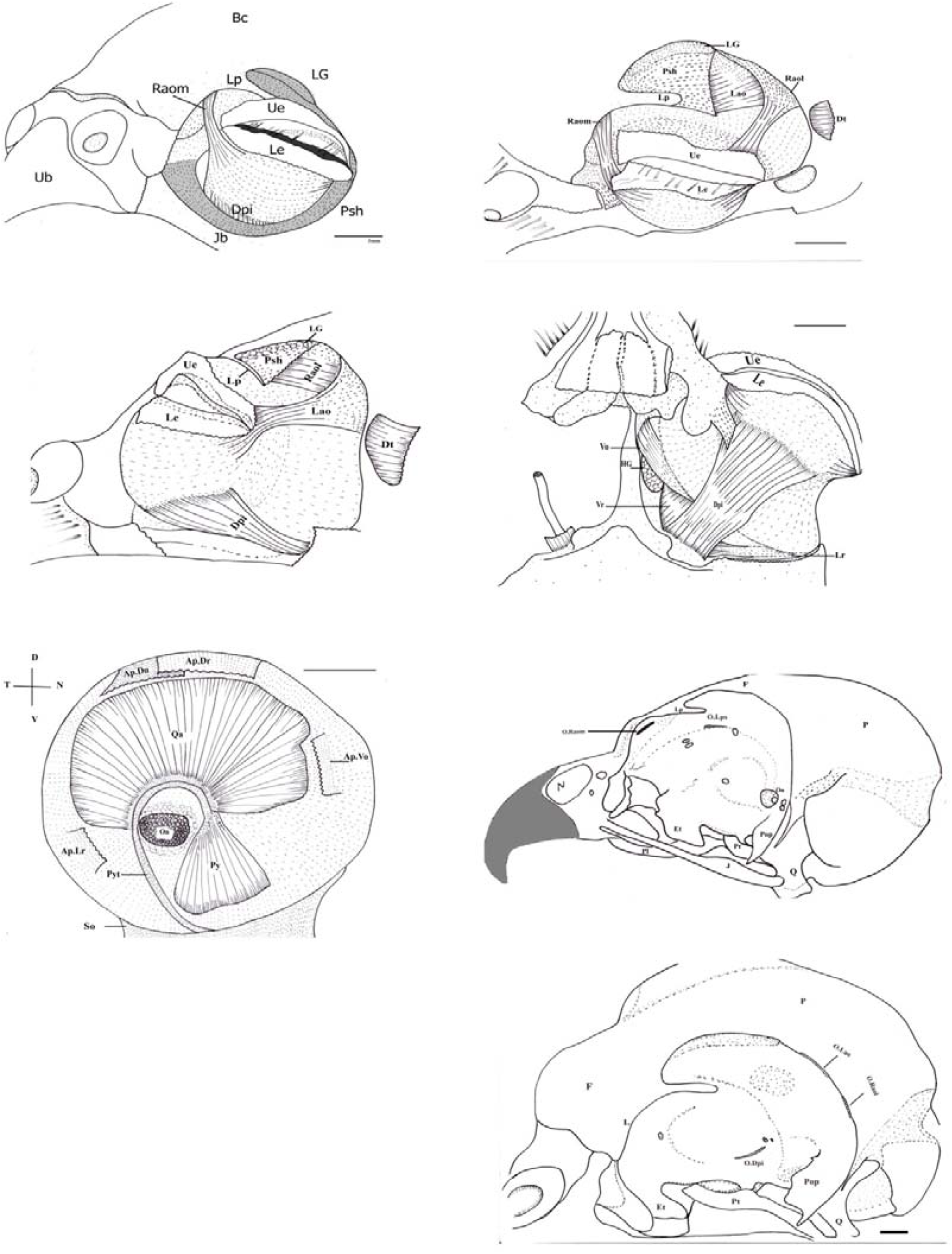
Photo of dissection of the eyelids muscles of little owl, *Athene noctua*, showing; the periorbital sheet (Psh), lacrimal process (Lp), the lacrimal gland (LG) Retractor anguli oculi Lateralis muscle (Raol), Retractor anguli oculi medialis muscle (Raom), Levator anguli oculi muscle (lao), dermato temporalis muscle (Dt), lateral rectus muscle (Lr), aponeurosis of lateral rectus muscle (Ap.Lr), ventral rectus muscle (Vr) and aponeurosis of ventral oblique muscle (Ap.Vo) which covers the harderian gland (HG), quadratus muscle (Qa), prymidalis muscle (py), prymidalis tendon (pyt), aponeurosis of dorsal rectus muscle (Ap.Dr), aponeurosis of dorsal oblique muscle (Ap. Do), the origin site of Levator palpebral superioris muscle (O.Lps) and depressor palpebral inferioris muscle (O.Dpi), Retractor anguli oculi Lateralis muscle (O.Raol), Retractor anguli oculi medialis muscle (O.Raom).

**Fig. 8.**
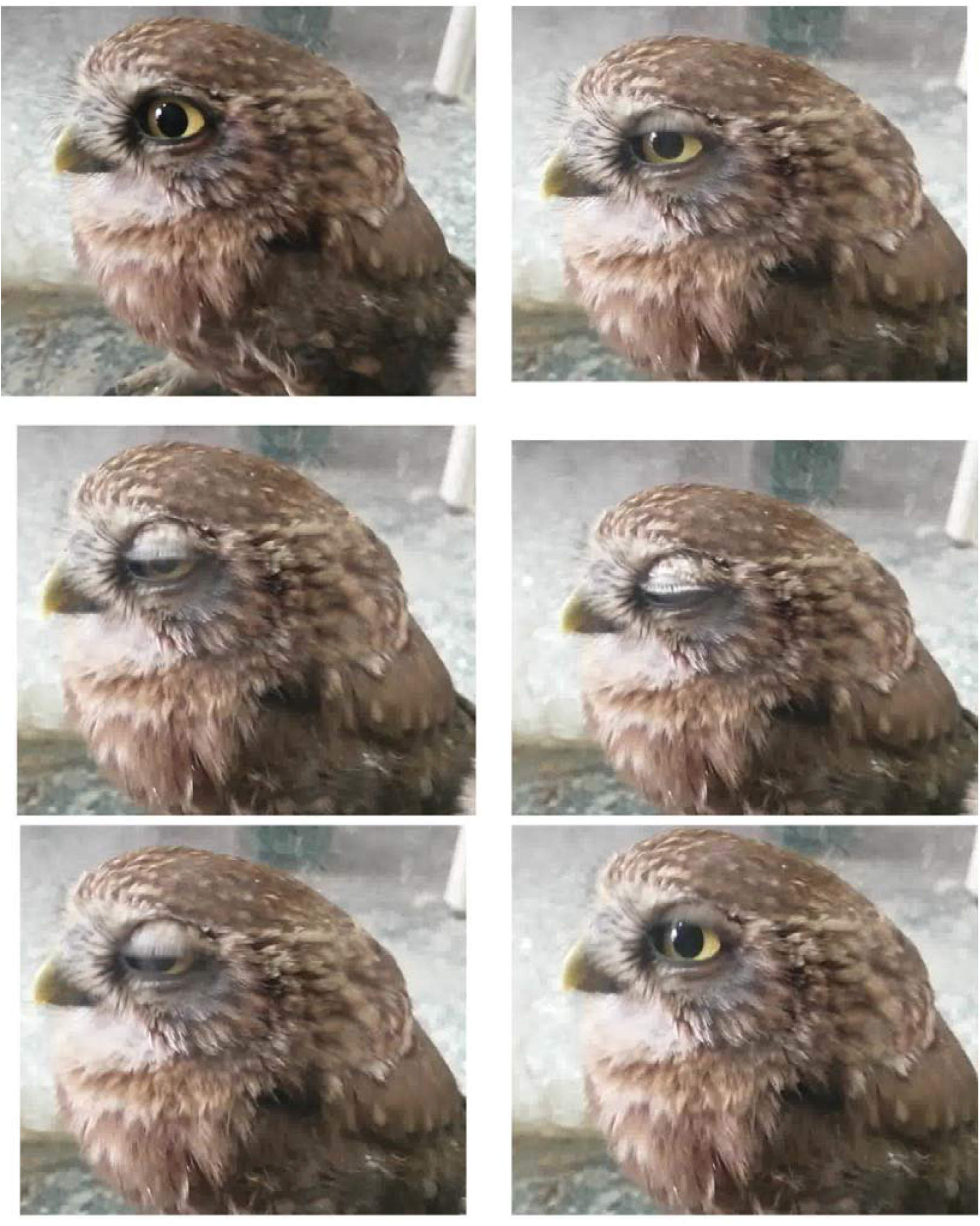
Photo of the sequent video record of movement of eyelids of little owl, *Athene noctua*, in natural condition.

**Fig. 9.**
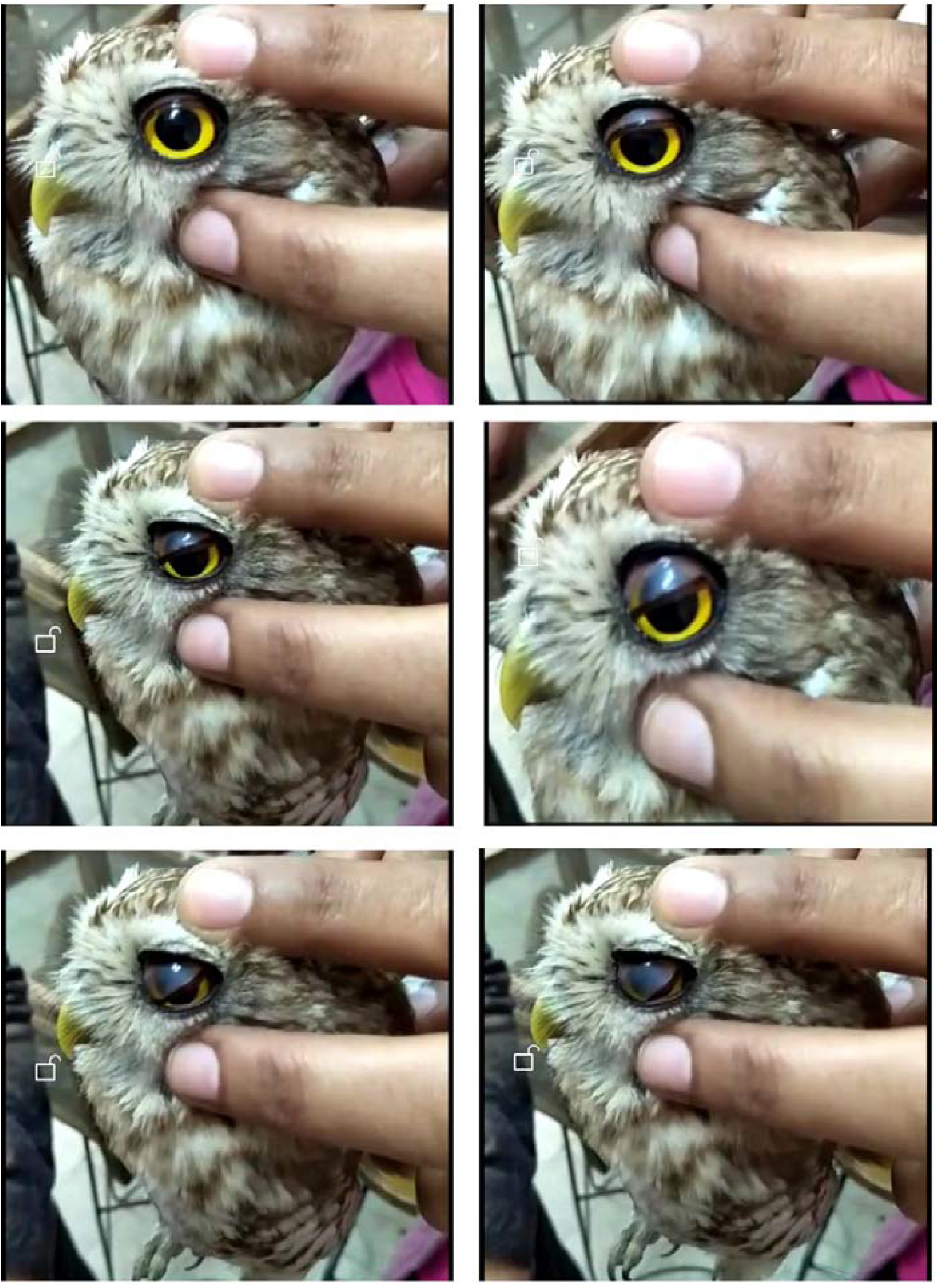
Photo of sequent of video record of movement of the nictitating membrane of little owl, *Athene noctua*, showing this membrane slips rapidly and obliquely across the front of eye from fixed pointed towards ventrotemporal direction

### Levator palpebral superioris muscle (Lps)

Levator palpebral superioris is sheet like with parallel muscle fibers that runs dorsal to the rectus dorsalis muscle. The Levator palpebral superioris extends dorso-laterally from its origin on the interorbital septum posterior to the origin site of the oblique dorsalis muscle. The muscle fibers of this muscle fibers fan out laterally to be inserted on the connective tissue of the upper eyelid. During its diverged, the muscle fibers interfere with the antero-dorsal fibers of the muscle Levator anguli oculi and under pass the glandula lacrimalis hides completely by the periorbital sheet which locates on the interorbital septum just then these muscles and interfere with and during its dorsolateral diverge the muscles of the Levator palpebral superioris.

### Levator anguli oculi muscle (Lao)

Levator anguli oculi is a sheet like, parallel fibred muscle which is located posterior-lateral to the muscle levator palpebral superioris and anterior to the muscle levator anguli oculi lateralis. The muscle levator anguli oculi originates from the posterio-lateral edge of the parietal bone (anterior temporal crest) via thin aponeurosis and fleshy muscle fibers, just anterior to the origin site of the muscle retractor anguli lateralis. Then the muscle levator anguli oculi runs obliquely antero-lateral to be inserted on the connective tissue of the upper eyelid and nictitating membrane and emerge with the muscle fibers of the Levator Palpebral Superioris.

### Retractor Anguli Oculi Medialis muscle (Raom)

Retractor anguli oculi medialis is a ribbon-like and parallel fibred muscle extends laterally from the anterior angle of the brain case to be run ventrally reaching the anterior angle of the eyelid. Along the extension of this muscle over pass the nasolacrimal sac. The muscle retractor anguli oculi medialis originates from the medial surface of the lacrimal bone then extends antero-ventral towards its insert. The insertion of the retractor anguli oculi medialis is diverged; some fibers attach on the anterodorsal surface of the upper palpebral margin while other fibers reach to the lower eyelids and spread fans out on the inferior tarsal plate.

### Retractor Anguli Oculi Lateralis muscle(Raol)

Retractor anguli oculi Lateralis is sheet like with parallel-fibred muscle extends from the posterior angle of the brain case to be attaching on the posterior angle of the eyelids. The muscle retractor anguli oculi lateralis originates from the posterior rim of the orbit (anterior temporal crest) via fleshy fibers and aponeurosis which over passes and covered partially the dorsal surface of the muscle Levator anguli oculi. The origin site of the muscle retractor anguli oculi lateralis locates posterior and anterolateral to the origin site of levator anguli oculi and dermotemporalis muscle respectively. The insertion of the retractor anguli oculi lateralis attaches on the posterior angle of the upper eyelid and few muscle fibers diverges to attach on the posterolateral surface of the inferior tarsus plate.

### Depressor Palpebral Inferioris muscle (Dpi)

The muscles depressor palpebral inferioris is fan-shaped like and parallel fibred muscle lies dorsal to the muscle obliqus lateralis and rectus ventralis. The ventral surface of this muscle is covered partially by the periorbital sheet. The depressor palpebral inferioris extends from the origin that lies on the interorbital septum (orbitosphenoid) just ventral to the optic foramen, medial to basiopterygoid process then fan out towards its insertion that locates on the posterior margin of the inferior tarsal plate.

### Quadrates muscle (Qa)

The quadrates muscle is broad, sheet and fan-like muscle located transversally dorsal to the optic nerve spread over almost the dorsal surface of the posterior segment of the eye ball. The quadrates muscle originate from the postero-dorsal edge of the sclera cartilage ventral to the insertion site of the rectus dorsalis and oblique dorsalis muscles. The muscle fibers extended posteriorly to insert via thin aponeurosis is on the tendon of the pyramidalis.

### Pyramidalis muscle (Py)

The muscle pyramidalis is a triangle shape with parallel fibers, lies ventral to the rectus ventralis muscle. The muscle pyramidalis is located on the anteroventral surface of the sclera cartilage, and then extends posteriorly towards the optic nerve as a tendon. This tendon runs postero-ventrally then antero-dorsally toward the nictitating membrane to attach on the anterior edge of its free margin.

## Discussion

The eyelids are unique protection tissues which extend from the skin to cover the eye socket when required. The eyelid movement is made involuntarily to protect the eye from the strong light or any foreign bodies, as well as, keep the cornea surface moist (Jochems and Phillips, 2015). The presence of three eyelids; upper, lower, and third eyelid which is also known as the nictitating membrane is common feature in birds (Baumel et al., 1993). Together with the orbital glands, the eyelids supply the moisture of the eye and maintain their health (Kleckowska-Nawrot et al., 2016).

The movement of the eyelids of the little owl was recorded in Lab and observed the upper eyelid moves rapidly and frequently down toward the lower eyelid, while the slipping of the nictitating membrane occur when fixed the movement of eyelids. The nictitating membrane slips in ventrotemporal direction. The movement of upper eyelid in little owl is opposite the views of most previous authors who’s confirmed the lower eyelid of bird is more mobile than upper one (Wyneken, 2012). That feature is common in mammals; the upper eyelid is more mobile and larger than the lower eyelid (Klećkowska-Nawrot et al., 2019).

The present authors supposed that the little owl is one of owl species hunts during the day, waiting and focusing on their prey then capture them by sharp claws (Mikkola and Heimo, 2013). The nature of the protective apparatus may be expected to vary according its activity pattern. The little owl being mostly less near the ground, thereby less exposed to dust and dirt. It is may expected that this owl is not needed to special mechanism to remove foreign bodies. The nictitation is happens to cleanse the front of the cornea but in little owl, this function performed by upper eyelid. Chauveau-Arloing (2018) pointed out the size of the nictitating membrane is inversely proportional to the ability of the animal to remove foreign bodies from its eyes

Jochems and Phillips (2015) said that the eyelid movement is made involuntarily as a reflect action against any foreign bodies. The present authors agree with this view but suggested there are some conditions may stimulate the movement of eyelids. The anatomical results showed that the whole eye ball with their muscles is surrounding by collagenous periorbital sheet. The variations in the thickness of the periorbital sheet and the difference in its connections with the skull among vertebrates may reflect how much the importance of this sheet.it was noticed that there is no available information about the importance of the periorbital sheet.

In little owl, this periorbital sheet lies directly beneath the epithelium of eyelids and is covered the levator and depressor muscles that control the movement of upper and lower eyelid respectively. Furthermore, this sheet in little owl connects with some movable skeletal parts of the skull, so any change in the movement of these skeletal parts during the feeding process yield to pull the periorbital sheet. Physically, the morphological features and topographical position of the periorbital sheet of the little owl allow making a pressure force on the levator and depressor eyelids muscles that may stimulate the contraction of these muscles, thereby causing in closing or opening of eyelids. Theoretically, we concluded that the eyelid movement not only depends on the periorbital sheet but also the movement of the skeletal elements of the skull during the feeding system. Ostheim et al. (2020) studies the eyelid squinting during food pecking in pigeons. It was confirmed that the feeding and sensory systems are integrated systems. The present authors suggest that the periorbital sheet may play an important role in coordinating the movement of eyelids.

Moreover, during anatomical investigation of the eye of little owl observed the orbital region (and the various glands there in) is connected with the nasal region nasolacrimal duct (tear duct). This duct appears in the little owl at anterior in oblique position, near the nictitating membrane. The retractor anguli oculi medialis muscle passess over this duct during its ventral extension to attach on the lower eyelid. The retractor anguli oculi medialis muscle in the little owl is considered as a one of accessory eye muscles that sharing in closing the palpreal fissure by pull the anterior angle of lower eyelid. The role of this muscle was mentioned by Slonaker (1918) in English sparrow, Burk (1893) in pigeon and Stibbe (1928) in mammals.

According to the anatomical feature of this muscle in little owl, the present authors expected that the contraction of the retractor anguli oculi medialis muscle may making pressure on the nasolacrimal duct that may help it to perform its function in the draining the orbital fluid.

However, the histological structure of eyelids facilitates its sweeping motion without any injury. The present study reveals that the structure of two eyelids is very similar in the little owl. Histologically, the clear variation between the two eyelids noticed among the number of cell layers forming the epithelium of skin and palpebral surfaces of the eyelids, the degree of keratinization over their surface, as well as, the distribution of pigment granules within their cells. The skin surface of both eyelids of little owl is composed of keratinized squamous epithelium. The cell-layered of the skin surface is varied from one to two layers in lower eyelid while it reached approximately three to four layers in upper eyelid. Thus, the lower eyelid in little owl appears very thin more than upper lid. The keratin layers that cover their eyelids act as a protective armor. Kardong (2009) said that the keratinization occur where friction or direct mechanical abrasion insult the epithelium.

The present investigation has observed that the margins of eyelids and its conjunctiva surface are non-keratinized, except a small area of the conjunctiva surface of the lower eyelid, just near its palpreal margin. In 1979, Edward Moumenea has reported about presence of keratinization of the conjunctiva and he was assumed that the degree of keratinization probably does not depend on direct stimulation or initiation of keratin synthesis, but depends on maintenance of organized basal layer, as well as, the keratinized cells produce on an epithelial keratitis with subsequent vascularization of the cornea.

The surface of cells of the conjunctiva epithelium has carried a tiny microvilli “cytoplasmic extension” and contains mucous goblet cells. The goblet cells appear in the upper eyelid in large number in comparison to those in the lower one. A similar result has been described in Barred owl, *Strix varia* (Jochems and Phillips, 2015). It was found that these cytoplasmic extensions are seen also on the bulbar surface of the third eyelid. Schramm et al. (1994) pointed out that the densely clustered microvilli exhibit resorptive capacity toward substances with low molecular weights.

The histochemical analysis of the glands that scattered within the palpebral and bulbi conjunctiva showed that their secretions are mucous neutral and acid substances. These secretions distribute over the corneal surface by the movement of eyelids and aiding the tiny microvilli to protect the corneal surface from any foreign body and keep it moist. Furthermore, the acidic nature for these glands affords immunoprotection to the cornea.

These protective and cleaning functions are supplemented by the gliding movement of the nictitating membrane. The nictitating membrane is a highly-specialized neuro-muscular system (Stibbe, 1928). Previous studies have found that the nictitating membrane is appeared more prominent and well developed in the eye of some vertebrates e.g. birds and reptiles, as well as, many mammals which are exposed to wind, storm, dust, and sand (Schramm et al., 1994).

During the movement of the nictitating membrane crosses over the cornea surface may expose the cornea surface to great friction and resulting in loss of optical quality. The anterior corneal surface of the little owl is covered by a multilayer of cuboidal cells that proliferate superficially to form a disc of flattened cells. This type of cornea epithelium may counteract the abrasive forces which occur in this region during the sweeping process of this membrane. Simultaneously, the fluid that secretes from the orbital glands moistens this membrane and helps to decrease the abrasive force and facilitate its sweeping motion without any injury.

## Acknowledgements

The authors are thankful to Faculty of Science, Assiut University, Egypt, for providing the opportunity to conduct this work and electron microscopy units for assistance for SEM investigations

## Data availability statement

The data that support the findings of this study are available from the corresponding author upon request.

